# Absolute measures of time-difference-of-arrival positioning error in underwater acoustic telemetry setups

**DOI:** 10.64898/2026.08.24.746702

**Authors:** James Adam Campbell, Petter Lundberg, Franz Hölker

## Abstract

This brief communication presents two solutions for calculating absolute measures of error from time-difference-of-arrival (TDOA) positioning in underwater acoustic telemetry arrays. First, a Monte Carlo estimation of TDOA positioning error is derived. Next, a computationally inexpensive, approximate solution to the Monte Carlo method is presented. This approximate solution is achieved by solving the Jacobian of a closed-form TDOA positioning model. The positioning error covariance matrix returned from either method can then be used to report the accuracy of TDOA positions or utilized in state-space positioning models. Finally, calculations of the expected radial error are shown which serves as a simple summary statistic for reporting positioning error in real units.

## 2 Introduction

Acoustic telemetry arrays are widely used to track the movements of tagged aquatic animals *in situ* (reviewed in Lennox et al. 2023). When telemetry receivers are placed closely together with overlapping detection ranges, time-difference-of-arrival (TDOA) positioning algorithms can estimate the locations of these tag emission events. These resulting location estimates can then be utilized in a variety of animal movement analyses (see Hooten et al. 2017).

Despite the prevalence of TDOA positioning in telemetry studies, the practice of reporting or utilizing absolute measures of positioning error appears to be uncommon. Given that many commercially available receivers come equipped with acoustic tags embedded within them—often termed sync-tags—positioning error can be readily calculated in many array setups. These sync-tags can be used to synchronize the clocks of independent receivers in an array and, importantly, serve as an empirical measure of detection error. Here, detection error is the time difference between the measured and expected detection times for a tag transmission arriving at a receiver.

In the following sections, two methods for transforming these empirical measures of detection error into absolute positioning error covariance matrices are demonstrated. Finally, practical solutions for reporting positioning error are shown by (i) plotting 95% confidence ellipses around location estimates and (ii) the calculation of the expected radial error along with its variance.

## 3 Simulated positioning error

Assuming the detection error distribution is Gaussian with a known scale parameter, *σ*_det_ (measured from sync-tag detections), the positioning error at a particular location, **x** = (*x*_*x*_, *x*_*y*_), can be estimated via Monte Carlo simulation with

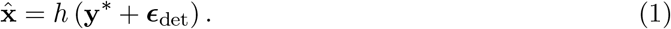

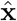 is an estimate of the true tag location, **x**, returned by the TDOA positioning function, *h*(), given the expected detection times 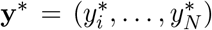 across *N* receivers with additional randomly sampled detection error

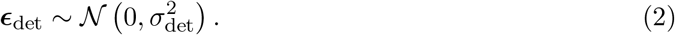

The expected detection times for a tag at location **x** are given by

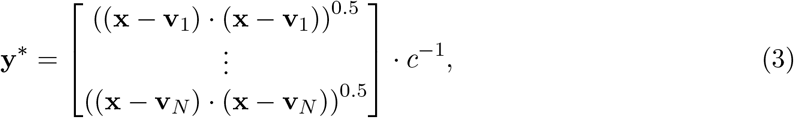

where **v**_*i*_ is the location of receiver *i* and *c* is the transmission speed.

For a given location, **x**, many instances of 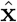 can be simulated. The covariance matrix of these estimates can then be taken as a simple measure of positioning accuracy. This can be plotted as a 95% confidence ellipse over a map of the study area or used to compute a summary statistic for positioning accuracy.

## 4 Closed-form positioning error

In the case that *h*() is a differentiable function, the positioning error covariance matrix can be approximated by

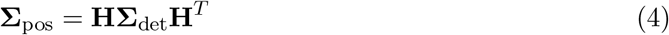

and

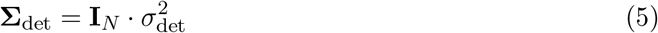

where **H** is the Jacobian of *h*() and **I**_*N*_ is a *N* by *N* identity matrix. In this section, the solution of **H** will be shown for Smith and Abel‘s (1987) closed-form TDOA method, termed *spherical interpolation*.

For an array of receivers arranged in *D* dimensional space, where there are at least *D* + 2 detections, TDOA position estimates can be given by

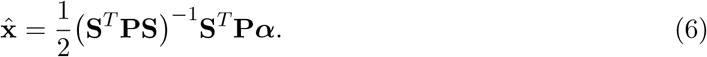

**P** is a projection matrix defined by

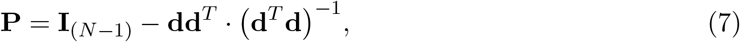

where **I**_(*N* −1)_ is a (*N* − 1) by (*N* − 1) identity matrix. The matrix **S** holds the receiver locations relative to a chosen reference receiver—**v**_1_ in this case—where

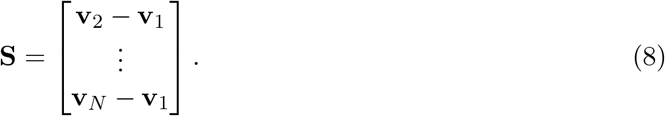

The remaining terms are given by

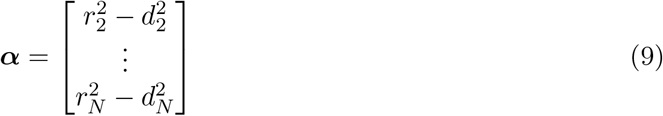

and

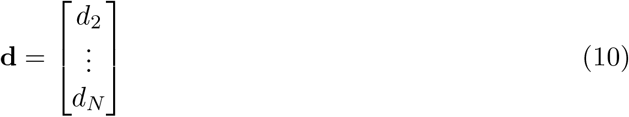

where *r*_*i*_ is the distance between receivers *i* and 1. *d*_*i*_ is the distance difference of the source between receivers 1 and *i*, estimated by

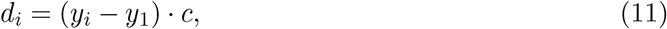

where *y*_*i*_ is the measured detection time at receiver *i*. The above formulation is a simplified implementation of Smith and Abel‘s (1987) *spherical interpolation* solution where the weighting matrices have been set to an identity matrix.

The solution for the Jacobian with respect to the measured detection times, **y**, is then given by

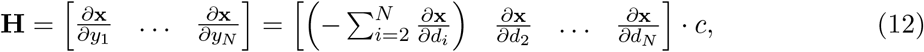

where

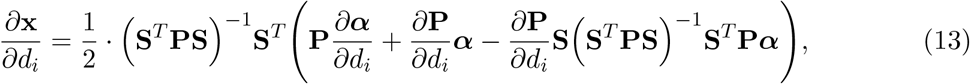

and

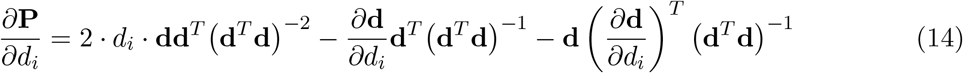

with

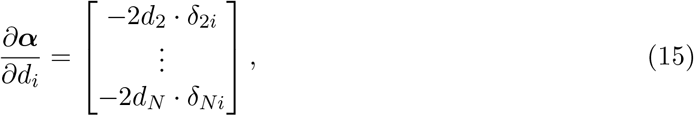

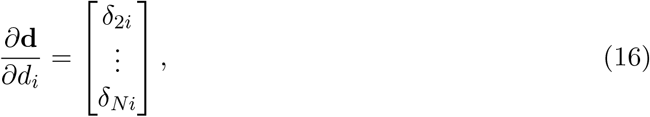

where *δ*_*ji*_ is the Kronecker delta function.

The Jacobian, **H**, can be calculated with either (i) the expected detection times from a known tag location, **y**^∗^, or (ii) the measured detection times from an unknown tag location, **y**. Figure 1a shows example 95% confidence ellipses calculated for Smith and Abel‘s (1987) TDOA solution. Both the Monte Carlo and Jacobian methods have been used to estimate the positioning error covariance. Here, the confidence ellipses were calculated using the known locations of four simulated tags in a small telemetry array of 6 receivers. The detection error was set to *σ*_det_ = 1 ms. The Monte Carlo method used 1000 simulated positions for each confidence ellipse.

**Figure 1:**
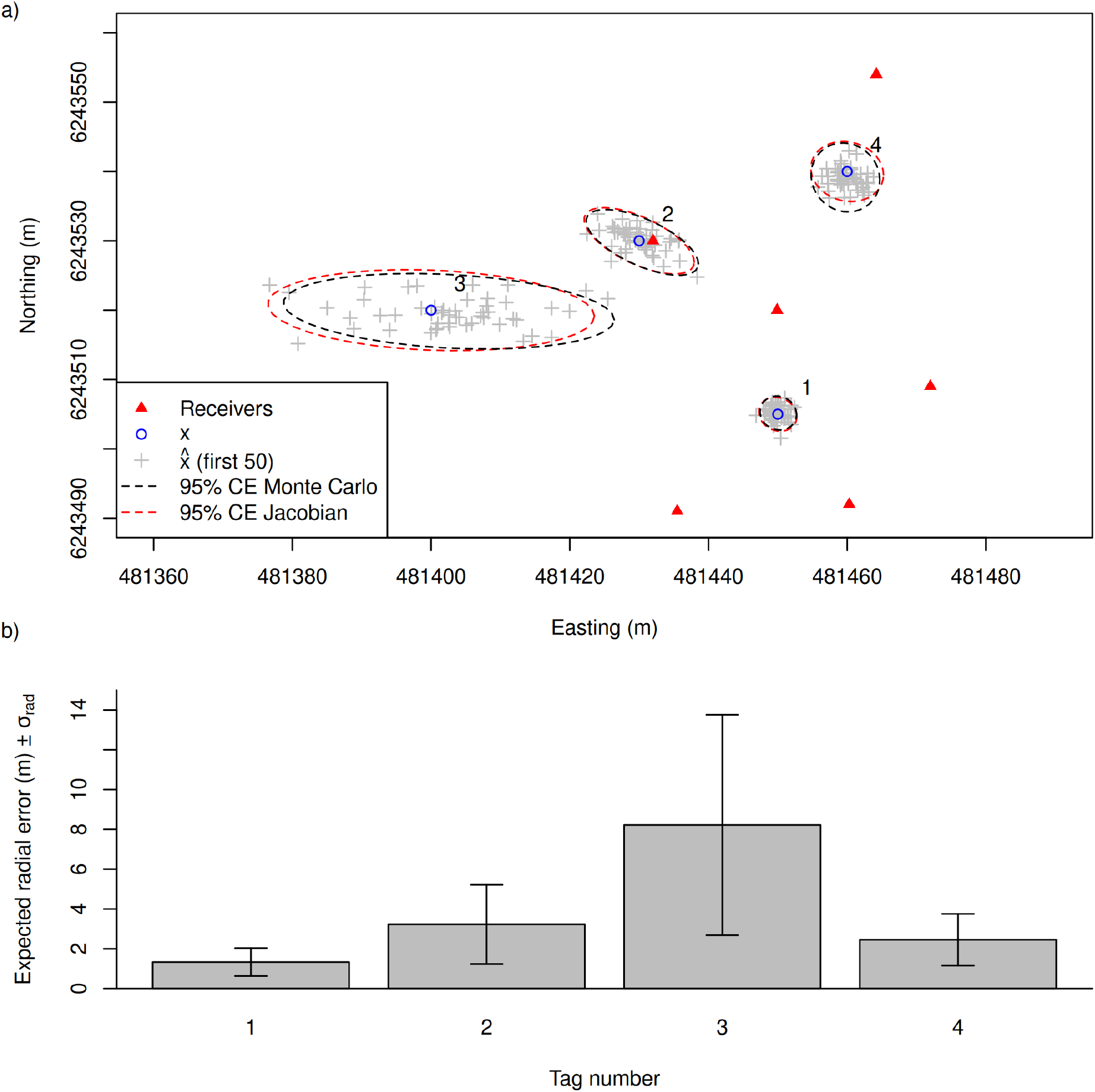
Panel (a) shows an example array with six receivers where positioning error is estimated at four locations, **x**, with a detection error scale parameter set to *σ*_det_ = 1 ms. For each location, a 95% confidence ellipse has been generated for Smith and Abel‘s (1987) closed-form TDOA solution using both the Monte Carlo (1000 simulations) and Jacobian methods. The Jacobian method is extremely fast to execute and bears a close resemblance to the Monte Carlo simulated ellipses. Panel (b) shows the expected radial error for each of the 4 locations calculated using the Jacobian method.

## 5 Reporting positioning error

After the positioning error covariance is calculated, it can be visualized by plotting 95% confidence ellipses around the location—or an estimate of the location. Here, the Cholesky decomposition of the positioning error covariance matrix can be used to transform a scaled circle into a confidence ellipse

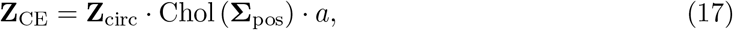

where **Z**_circ_ is a matrix of coordinates for a unit circle centered around (0, 0), Chol (**Σ**_pos_) is the upper triangular factor of the Cholesky decomposition of the positioning error covariance matrix, and *a* is a scaling term. For a 95% confidence ellipse, 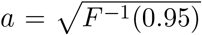, where *F* ^−1^ is the inverse of the Chi squared cumulative distribution function with two degrees of freedom. The resulting confidence ellipse, **Z**_CE_, can then be centered on the tag location as demonstrated in figure 1a.

Additionally, the error covariance can be concisely reported with a simple summary statistic: the expected radial error. That is, the expected distance in meters between the true and estimated tag location. A solution for this is provided by Chew and Boyce (1962) where

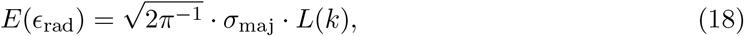

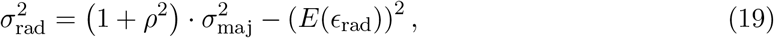

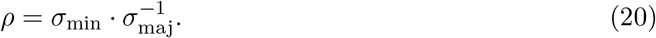

Here, *ϵ*_rad_ is the radial error in meters, where *E*(*ϵ*_rad_) and 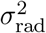 are then the expected value and variance of the error. *σ*_min_ and *σ*_maj_ are the standard deviation from the minor and major axes of **Σ**_pos_, respectively, *ρ* is the ratio of those standard deviations, and *L*(*k*) is the elliptic integral of the second kind with respect to the eccentricity, 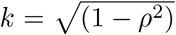. Please refer to the original paper for more details on this formulation.

Figure 1b shows the radial error estimated for each of the four locations in the example array. Here, the Jacobian method was used to estimate the positioning error covariance matrix, **Σ**_pos_, from the expected detection times from the known locations and then singular value decomposition was used to retrieve *σ*_maj_ and *σ*_min_,

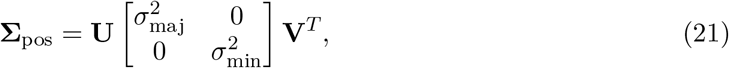

where the variance along the minor and major axis, 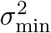 and 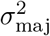, are the singular values and the columns of **U** and **V** hold the left and right singular vectors.

## 6 Expected versus measured detection times

The methods described in this manuscript work well for reporting positioning error when the location of the tag and the expected detection times are known beforehand. In practice, however, researchers are often interested in positioning error when the location of a tagged animal is not known. These measures can be used to, among other purposes, (i) filter out position estimates with low accuracy, or (ii) apply state-space positioning models which take that measurement error into account (see Campbell et al. 2025). In this case, only the measured detection times are available to calculate the positioning error.

Campbell et al. (2025, figure 4b) showed when equation (12) is calculated with measured detection times, reasonably good measures of positioning uncertainty are still returned—granted no large outliers were present. However, in the presence of large outliers resulting from reflected detections, the positioning error tended to be greatly underestimated. Hence, these methods should not be applied to datasets where reflected transmissions are expected to be prevalent.

Figure 2 shows the effect of increasing detection error on the resulting estimates of radial error when using measured detection times. Here, the standardized residuals of the radial error are plotted for simulated tag emissions from each of the four locations in the example array. At each location, 1000 emissions have been simulated for the following scenarios, *σ*_det_ = 0.1, 1, 2 and 3 ms. Here, the expected radial error was estimated from the measured detection times using the Jacobian method—that is, without information on the true tag location. The true radial error (distance between the true and estimated tag location) was then subtracted by the expected radial error and then standardized by the standard deviation, *σ*_rad_. If the positioning error is estimated well, the resulting residuals will fall along a standard normal distribution.

**Figure 2:**
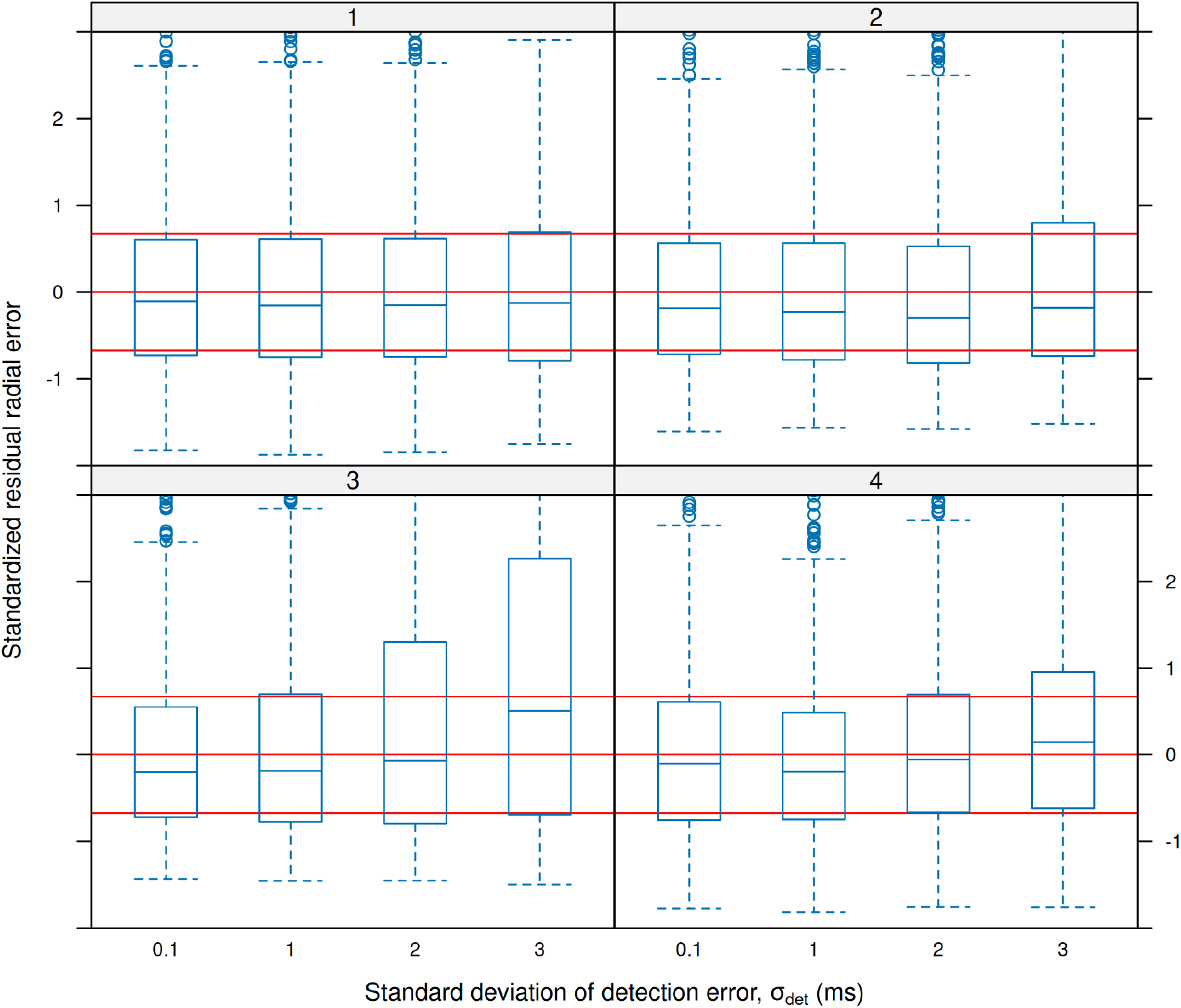
Standardised residuals of the radial error at each of the four locations in the example array (marked in the facet labels). At each location, 1000 tag emissions were simulated for each *σ*_det_ = 0.1, 1, 2 and 3 ms, where the radial error was then estimated from the measured detection times. When the radial error is estimated well, the resulting residuals will be standard normally distributed—i.e. the box plot quartiles will align well with the red annotations.

As seen in figure 2, even when using measured detection times with large error, the residual radial error is reasonably well distributed for those locations within the array. For the location outside the array, increasing detection error resulted in overdispersed residuals—the estimated radial errors matched poorly with the true simulated values. Note that this figure is provided as a proof of concept as it represents just a few tag locations where each emission is detected by all six receivers in the example array. In practice, researchers should make use of more detailed simulated datasets to verify that the resulting radial error estimates are suitable for their particular telemetry setups.

## 7 Conclusion

In this brief communication, a simple and fast method for computing the covariance matrix of absolute positioning error has been demonstrated by solving the Jacobian of a closed-form TDOA positioning solution. This approximate solution was shown to perform well when compared with error estimates calculated from Monte Carlo simulations. Furthermore, these error covariance matrices were then used to calculate the expected value and standard deviation of the radial error; providing a simple summary statistic for reporting positioning error in real units (meters). Even when the true location of the tag is not known and only the measured detection times are available, reasonable estimates of positioning error can still be provided—ideally paired with some verification procedure using simulated data.

In the described methods, the detection error is assumed to be Gaussian with a scale parameter that is known beforehand. As many telemetry arrays come equipped with sync-tags, measuring the detection error distribution can be easily done.

Lastly, example code has been provided as supplementary material: An R implementation of the described methods along with a reproducible document exemplifying their use.

## Supporting information

supmat_functions.R

Example R implementation

## 8 Acknowledgements

This project has received funding from the following: European Union Horizon 2020 Research and Innovation Programme under the Marie Sklodowska-Curie Actions, grant agreement no. 860800.

## 9 Declaration of Generative AI and AI-assisted technologies in the writing process

Generative AI or AI-assisted technologies were not used at any stage in the preparation of this manuscript or the work presented in the manuscript.

