## Supplementary material for "Absolute measures of time-difference-of-arrival positioning error in underwater acoustic telemetry setups": supmat_functions.R: supmat_functions.R

```
# Smith and Abel's TDOA positioning (Spherical Interpolation)
#
# This is a closed-form solution for TDOA positioning, termed spherical
# interpolation.  It's a good approximation to the maximum likelihood solution
# and can be applied when there are at least (D + 2) detections, where D is the
# number of spatial dimensions.
#
# Parameters
#  - y: A vector of detection times.
#  - receivers: A matrix holding the Easting/Northing coordinates of each
#     receiver in the array.
# - c:  Speed of signal transmission (m/s).
#
func_h <- function(y, receivers, c){

  receivers <- as.matrix(receivers)

  y <- as.vector(y)

  # Remove NA values
  idx_keep <- which(!is.na(y))
  y <- y[idx_keep]
  receivers <- receivers[idx_keep,]

  # Number of receivers
  N <- nrow(receivers)

  # Coords of receivers, relative to the first one.
  mS <- apply(receivers[-1, ], 1, `-`, y = receivers[1,], simplify = F) |>
    do.call(what = "rbind") |> as.matrix()
  # Vector of distance differences (relative to first receiver)
  vd <- c*(y[-1] - y[1])
  # Distance between receivers and reference
  vr <- sqrt(mS[,1]^2 + mS[,2]^2)

  valpha <- vr^2 - vd^2
  mP <- {
    mI <- diag(rep.int(1, times = length(vd)))
    mI - ((vd %*% t(vd)) / ((t(vd) %*% vd)[1]))
  }
  mPP <- mP %*% mP
  # Eq. 14, Smith & Abel (1978)
  vx <- 0.5 * solve(t(mS) %*% mPP %*% mS) %*% (t(mS) %*% mPP %*% valpha)

  return(as.vector(vx + receivers[1,]))

}

# Calculate expected detection times
#
# For a given location, this will return the expected detection times.
#
# Parameters
#  - x: The coordinates of a tag.
#  - receivers: A matrix holding the coordinates of the receivers in the array.
#  - c: Speed of signal transmission.
#
func_toa <- function(x, receivers, c){
  x = as.vector(x)
  vg <- apply(receivers, 1, FUN = \(v) sqrt((x - v) %*% (x - v))[[1]] )
  return(vg/c)
}

# Calculate the Jacobian of Smith & Abel's TDOA solution
#
# This is the closed-form solution for the Jacobian of Smith & Abel's spherical
# interpolation TDOA positioning method.  This can be used for transforming
# detection error covariance matrices into positioning error covariance
# matrices.
#
# This is a matrix of partial derivative solutions for the estimated tag
# position, x, with respect to the vector of measured detection times, y.
#
# Parameters
#  - y: A vector of detection times.
#  - receivers: A matrix holding the Easting/Northing coordinates of each
#     receiver in the array.
# - c:  Speed of signal transmission (m/s).
#
func_jacobian_x <- function(y, receivers, c){


  # Remove NA values
  idx <- which(!is.na(y))
  y = y[idx]
  receivers = receivers[idx,]

  if(length(y) < (ncol(receivers) + 2))
    stop("Too few detections.")

  # ------ Initialize constant variables
  # Init matrix S
  # Positions of remaining receivers, relative to the first
  mS <- apply(receivers[-1,], 1, \(v) v - receivers[1,]) |> t()

  # Get vector of distance differences
  vd = as.matrix((y[-1] - y[1]) * c)

  # Get vector r
  vr <- apply(mS, 1, \(x) sqrt(x %*% x)) |> as.matrix()

  # Get vector alpha
  valpha <- vr^2 - vd^2


  # Init matrix P
  mP <- {
    mI <- diag(rep.int(1, times = length(vd)))
    mI - ((vd %*% t(vd)) / ((t(vd) %*% vd)[1]))
  }

  # Calculate reused portions of equations
  mStPS_inv <- solve((t(mS) %*% mP %*% mS))

  # Init empty return matrix
  mat_out <- matrix(NaN, nrow = ncol(receivers), ncol = length(y))

  # Loop through column indices
  for (i in 2:nrow(receivers)){

    # ------ Initalize loop variables
    # partial derivative of vector d wrt d
    p_vd <- {
      ret <- numeric(length(vd))
      ret[i-1] = 1; as.matrix(ret)
    }

    # partial derivative of vector alpha wrt d
    p_valpha <- {
      ret <- numeric(length(valpha))
      ret[i-1] = -2*vd[i-1]; as.matrix(ret)
    }

    # Partial derivative of matrix P wrt d
    p_mP <- {
      dT_d_inv <- 1/(t(vd) %*% vd)[1]
      (
        2 * vd[i-1] * (vd %*% t(vd)) * dT_d_inv -
          p_vd %*% t(vd) -
          vd %*% t(p_vd)
      ) * dT_d_inv
    }

    # Partial derivative of x wrt d
    mat_out[,i] <- {
      0.5 * mStPS_inv %*% t(mS) %*%
        (
          mP %*% p_valpha +
            p_mP %*% valpha -
            p_mP %*% mS %*% mStPS_inv %*% t(mS) %*% mP %*% valpha
        )
    }
  }

  # Change derivative base from d to y.
  # First column is the sum of the remaining columns times -1
  # This is because the first column is our reference receiver
  mat_out[,1] <- -apply(mat_out[,-1], 1, sum)
  return(mat_out * c)

}

# Calculate the radial error from a bivariate covariance matrix
# Method is taken from Chew and Boyce (1962).
#
# Parameters
#  - Sigma: The covariance matrix for a bivariate normal distribution
#
# Return
#  - Returns the expected radial value along with its standard deviation.
#
func_radial_error <- function(Sigma){

  if(any(dim(Sigma) != c(2,2)))
    stop("Sigma must be a 2 by 2 covariance matrix of positioning error.")

  # Apply Singular valued decomposition to covariance matrix
  # We have to rotate the matrix to remove the covariance terms from the
  # bivariate normal distribution
  svd <- svd(Sigma)

  # Standard deviation of the smaller orthogonal axis
  sig_1 <- sqrt(svd$d[2])
  # Standard deviation of the larger orthogonal axis
  sig_2 <- sqrt(svd$d[1])

  # The below solution for radial error is taken from:
  #  - Chew, V., Boyce, R. 1962. Distribution of Radial Error in the Bivariate
  #     Elliptical Normal Distribution. Technometrics, 4(1), pp 138--140
  rho = sig_1 / sig_2
  k = sqrt(1 - rho^2)
  L_k <- {pracma::ellipke(m = k^2)}$e
  # L_k is Legendre's complete elliptical integral of the second kind wrt
  # the ellipticity k

  # Expected radial error
  E_r <- sqrt(2/pi) * sig_2 * L_k

  # Standard deviation of radial error
  sigma_r <- sqrt((1 + rho^2)*(sig_2^2) - E_r^2)

  return(list(E_r = E_r, sigma_r = sigma_r))

}
```
