## Supplementary material for "Absolute measures of time-difference-of-arrival positioning error in underwater acoustic telemetry setups": Example R implementation

Here we'll show how to apply the positioning error methods from the main manuscript within R. First, load the supplementary file. This contains R implementations of the equations described in the manuscript. Descriptions of these functions are provided above their definitions in the source file. Where possible, function and variable names have been chosen to match the notation used in the main manuscript.

```
source("./supmat_functions.R")
```

We'll make an example study area containing 6 telemetry receivers. We'll then mark four tag locations where we'd like to calculate the absolute positioning error for.

```
# Set receiver positions (Easting / Northing in meters)
receivers <- matrix(
  data = c(
    481460.3, 6243492,
    481435.5, 6243491,
    481449.9, 6243520,
    481464.2, 6243554,
    481432.0, 6243530,
    481472.0, 6243509),
  ncol = 2, byrow = T)

# Set four example tag locations
x <- matrix(
  data = c(
    481450, 6243505,
    481430, 6243530,
    481400, 6243520,
    481460, 6243540),
  ncol = 2, byrow = T)

# Set tag transmission speed
c <- 1500# m/s

# ------ Plot tags within array
# Create a convenience function for plotting the study area
xlim = range(receivers[,1], x[,1])
ylim = range(receivers[,2], x[,2])
plot_base <- function(){
  plot(NULL, xlim = xlim, ylim = ylim,
  asp = 1, xlab = "Easting (m)", ylab = "Northing (m)")
  points(receivers, col = 'red', pch = 17)
  text(x = x[,1]+5, y = x[,2]+5, labels = 1:4, adj = c(1,1))}
# Plot study area
plot_base()
```

```r
# Plot tags
points(as.matrix(x), col = "blue", pch = 1)
# Show legend
legend("bottomleft",
  legend = c("Receivers", "Tags"),
  col = c("red", "blue"), pch = c(17, 1))
```

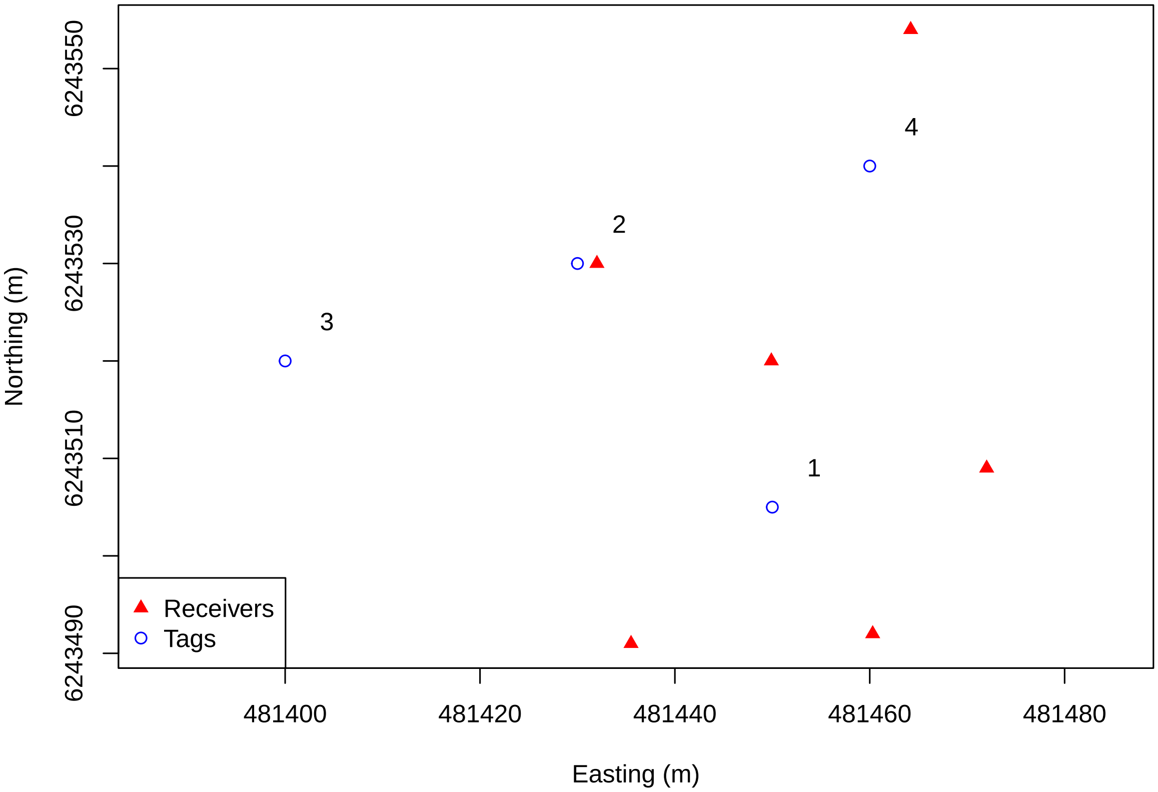


### Monte Carlo simulation

Here, we'll show how Monte Carlo simulation can be used to estimate the positioning error for a tag emission from a known location. First, we'll define a simple function for applying TDOA positioning to the expected detection times with some randomly sampled error added to it.

```r
# Create function for simulated erroneous position estimates
MC_positions <- function(x, receivers, sigma_det, n){

   # Init return matrix
  mat_ret <- matrix(NaN, nrow = n, ncol = ncol(receivers))

  # Get expected detection times
  y_star <- func_toa(
    x = x,#                 Coordinate of tag
    receivers = receivers,# Matrix of receiver coordinates
    c = c)# m/s             Speed of signal transmission

  # --- Loop until desired replications are reached
  for(i in 1:n){
```

```
    # Randomly sample detection error
    epsilon <- rnorm(n = nrow(receivers), sd = sigma_det)

    # Apply TDOA positioning to erroneous detection times
    mat_ret[i,] <- func_h(
      y = y_star + epsilon,# s Erroneous detection times
      receivers = receivers,#  Matrix of receiver coordinates
      c = c)# m/s              Speed of signal transmission
  }

  return(mat_ret)

}
```

We'll use this function to create a distribution of erroneous position estimates which we'll then measure the covariance matrix of. Let's apply this to one of our example tag locations.

```
# Standard deviation of Gaussian detection error
sigma_det <- 0.001# s

# Number of erroneous positions to simulate
n = 200

# Simulate erroneous positions
mX_hat <- MC_positions(
  x = x[2,],#                Coordinate of first tag
  receivers = receivers,#    Matrix of receiver coordinates
  sigma_det = sigma_det,# s SD of Gaussian detection error
  n = n)#                    Number of replications

# Measure covariance matrix from erroneous positions
mX_hat_Sigma <- var(mX_hat)

# Get mean value of erroneous positions
mX_hat_mu <- apply(mX_hat, 2, mean)
```

`mX_hat_Sigma` holds the positioning error covariance matrix for the second tag location. We can present this visually by creating a 95% error ellipse from it. Below is an example on how to do this via Cholesky decomposition.

```
# We'll make a convenience function for creating the confidence ellipses
conf_ellip <- function(Sigma, mu, p = 0.95){

  # Number of points to draw
  lngth = 40

  # Create unit circle
  circ_unit <- cbind(
    x = cos(seq(0,2*pi, length.out = lngth)),
    y = sin(seq(0,2*pi, length.out = lngth)))

  # Scale unit circle to desired percentile
  circ_conf <- circ_unit * sqrt(qchisq(p = p, df = 2))
  # A chi distribution with 2 degrees of freedom gives the distribution of
  # vector magnitudes for points sampled from a bivariate standard normal
```

```
  # distribution.

  # Now we'll use the Cholesky decomposition to scale our confidence circle
  # to match the covariance observed in our simulated positions.
  ellip_conf <- circ_conf %*% chol(Sigma)

  # Add the offset
  ellip_conf <- apply(ellip_conf, 1, `+`, mu) |> t()

  return(ellip_conf)

}
```

Below we've plotted an example confidence ellipse along with the erroneous position estimates.

```
# Plot study area
plot_base()

# --- Plot Monte-Carlo simulation results
# Position estimates
points(mX_hat, pch = 3, col = "darkgray")

# 95% confidence ellipse
lines(conf_ellip(mX_hat_Sigma, mu = mX_hat_mu, p = 0.95),
  col = "black", lty = 2)

# Mean position estimate
points(t(mX_hat_mu), col = "black", pch = 4)

# Plot true position
points(x[2,, drop = F], col = "blue", pch = 1)

# Show legend
legend(x = "bottomleft",
  legend =
    c("Receivers",  expression(x), expression(hat(x)),
      expression(mean(hat(x))), "95% CE"),
  col = c("red", "blue", "darkgray", "black", "black"),
  pch = c(17, 1, 3, 4, NA),
  lty = c(NA,NA,NA,NA,2))
```

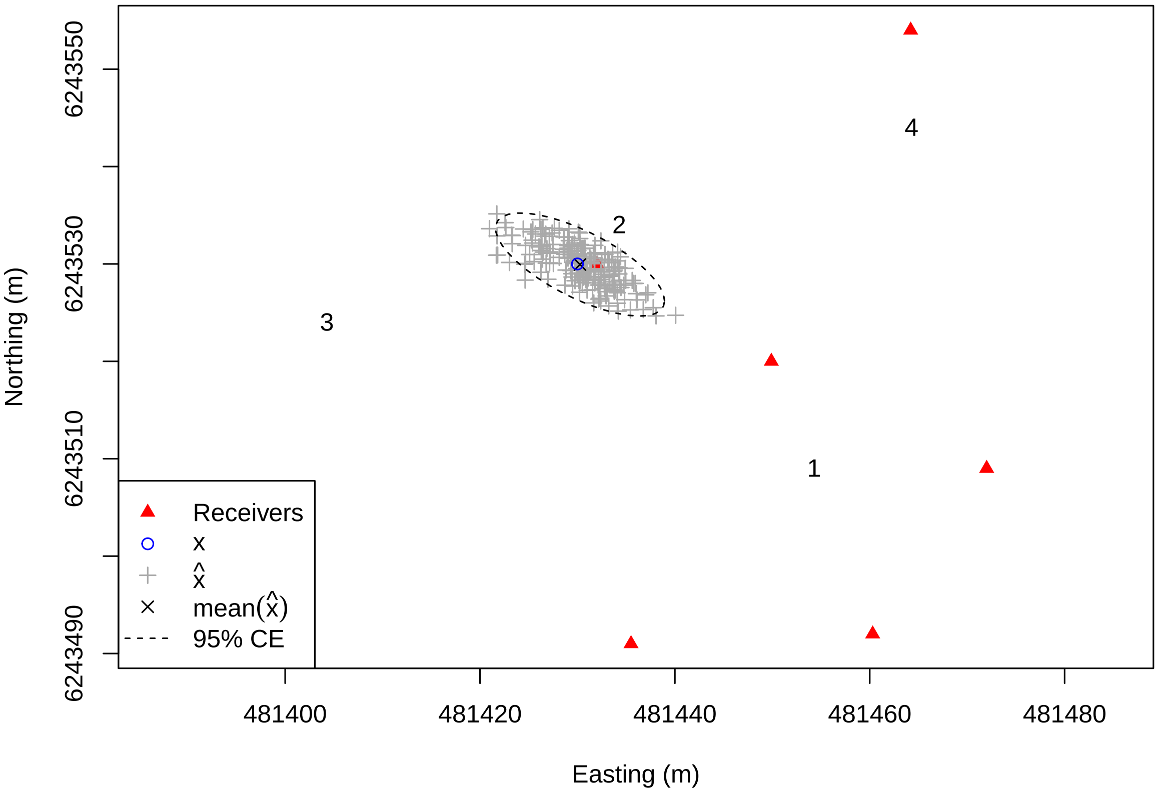


As you can see, the 95% confidence ellipse does a pretty good job of capturing the distribution of the positioning error. Monte Carlo simulations are handy for this purpose. However, accurate confidence ellipses (and covariance matrices) require many simulated positions. For large datasets with many locations, this can take significant computational resources.

### Closed-form positioning error

As described in the manuscript, we can instead calculate the required covariance matrix using the Jacobian of our TDOA function. This is much faster and provides a deterministic result. Below is a convenience function for calculating the closed-form positioning error covariance matrix. In this case, we're calculating the Jacobian from Smith & Abel's *spherical interpolation* solution for TDOA positioning.

```
get_Sigma_pos_closed <- function(x, receivers, c, sigma_det){

  # Get expected detection times
  y_star <- func_toa(
    x,#                     Coordinate of tag
    receivers = receivers,# Matrix of receiver coordinates
    c = c)# m/s             Speed of signal transmission

  # Get Jacobian of h()
  # This uses Smith & Abel's (1987) "spherical interpolation" TODA solution
  mH <- func_jacobian_x(
    y_star,#     Expected detection times
    receivers, # Matrix of receiver coordinates
    c)# m/s      Speed of signal transmission

  # Initialize detection error covariance
```

```
  mI <- diag(rep.int(1, times = length(y_star)))
  Sigma_det <-  mI * sigma_det^2

  # Transform into positioning error covariance
  Sigma_pos <- mH %*% Sigma_det %*% t(mH)

  return(Sigma_pos)
}
```

We'll plot the confidence ellipses for all the tag locations using both the Monte Carlo and closed-form methods below for a comparison.

```
# Plot study area
plot_base()

# Loop tag locations
for(i in 1:nrow(x)){

  # Get Monte Carlo positions
  pos_MC <- MC_positions(
    x = x[i,],
    receivers = receivers,
    sigma_det = sigma_det,
    n = 1000)
  # Plot first 50 erroneous position estimates
  points(pos_MC[1:50,], pch = 3, col = 'grey')

  # Get covariance matrix (closed-form)
  Sigma_pos_closed <- get_Sigma_pos_closed(
    x = x[i,],#               Tag coordinates
    receivers = receivers,#   Matrix of receiver coordinates
    c = c,# m/s               Speed of transmission
    sigma_det = sigma_det)# s SD of detection error

  # Plot closed-form confidence ellipse
  conf_ellip(Sigma_pos_closed, x[i,], p = 0.95) |>
    lines(col = "red", lty = 2)

  # Plot Monte Carlo confidence ellipse
  conf_ellip(var(pos_MC), apply(pos_MC, 2, mean), p = 0.95) |>
    lines(col = "black", lty = 2)

  # Plot tag location
  points(t(x[i,]), col = "blue", pch = 1)

}

# Draw legend
legend(x = "bottomleft",
  legend =
    c("Receivers", expression(x), expression(paste(hat(x),
      " (first 50)")), "95% CE Monte Carlo",
      "95% CE closed-form"),
  col = c("red", "blue", "gray","black", "red"),
```

```r
  pch = c(17, 1, 3, NA, NA),
  lty = c(NA,NA,NA,2,2))
```

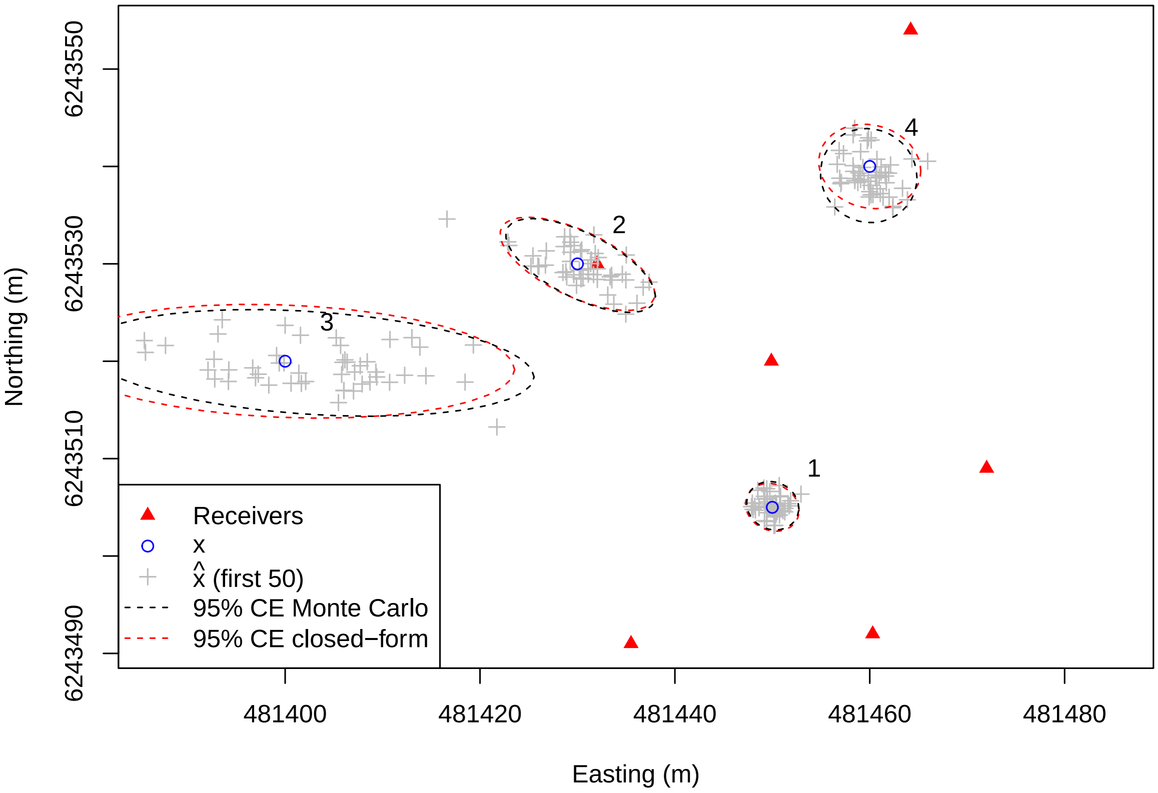


The closed-form solution matches up really well with our simulated results. Above, we increased our number of simulated points for each location to 1000 to get more stable and accurate estimates.

### Reporting positioning error

A simple measure of positioning error can be created by calculating the *radial error* from the position error covariance matrices. We've provided a function for this in the supplementary material.

```r
# Calculate radial positioning error for each location
df_r <- apply(x, 1, \(x_i){

  # --- Get Expected detection times
  y_star <- func_toa(
    x = x_i, #                Tag coordinate
    receivers = receivers, # Matrix of receiver coordinates
    c = c)# m/s               Speed of tag transmission

  # --- Calculate TDOA Jacobian
  H <-  func_jacobian_x(
    y_star,#      Expected detection times
    receivers, # Matrix of receiver coordinates
    c)# m/s      Speed of tag transmission

  # --- Transform detection error into positioning error
  Sigma_pos <- H %*% (diag(rep(1, times = 6))*sigma_det^2) %*% t(H)
```

```
  # --- Return radial positioning error
  return(as.data.frame(func_radial_error(Sigma_pos)))

}) |> do.call(what = 'rbind')

# Add tag labels
df_r$tag = factor(1:4)

# Plot of radial positioning error
barplot(E_r ~ tag, data = df_r,
  ylab = expression(paste("Expected radial error (m) \u00b1 ",
    sigma[rad])), xlab = "Tag number",
  ylim = c(0, 15)) |>
  arrows(y0 = df_r$E_r - df_r$sigma_r, y1 =  df_r$E_r + df_r$sigma_r,
    angle = 90, code = 3)
# Draw box around plot
box(bty = "l")
```

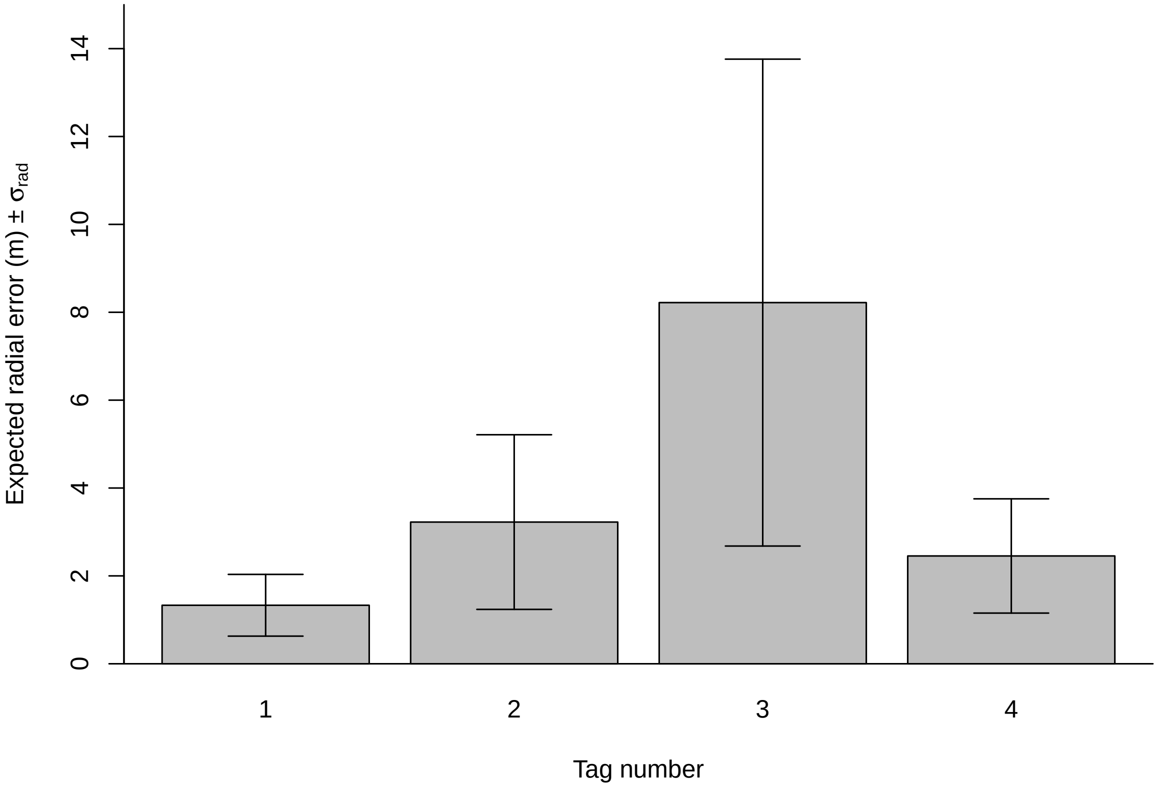


The above plot shows the expected (i.e. average) positioning error for each of the four locations along with the standard deviation of the radial error.

### Measured deteciton times

It's often the case that positioning error measures are desired when the expected detection times (along with the tag location) are unknown. When using measured detection times in place of the expected times, the resulting positioning error will be of a lower quality.

Here, it can be a good idea to use simulations to verify how closely the estimated positioning error resembles the true positioning error—for a given tag location and scale of detection error in your array. In the code below, we'll

1. simulate 1000 random emissions with detection error at each of our four examples locations for the following detection errors, $\sigma_{\text{det}} = 0.1$ ms, 1 ms, 2 ms, and 3 ms.
2. Apply the closed-form solution for estimating the positioning error covariance for measured detection times;
3. and then report the standardized residuals of the radial error estimates.

Here, the *radial error residuals* are the difference between the true radial error (distance between the true and TDOA estimated tag position) and the estimate of the expected radial error from the measured detection times. We can then standardize the residuals by dividing them by the estimated standard deviation of the radial error.

When the estimated radial error matches well with the true values, the resulting standardized residuals will fall along a standard normal distribution. This can act as a simple method to verify if radial error values resulting from measured detections are trustworthy for a given array setup.

```
# Set detection error values
sigma_dets <- c(0.1,1,2,3) * 1e-3

# Initiate dataframe for holding results
df_results <- data.frame()

# Loop all four tag locations
for(i in 1:4){
  # Loop all four detection error values
  for(j in 1:4){

    # Get detection error value
    sigma_det = sigma_dets[j]

    # Get true tag location
    x_i <- x[i,]

    # Get expected detection times for this location
    y_star <- func_toa(
      x = x_i,#                 Coordinate of true tag location
      receivers = receivers, # Matrix of receiver coordinates
      c = c)# m/s               Speed of transmission

    # Add simulated error to detection times
    Y <- replicate(
      n = 1000,# Number of simulations
      expr = {y_star + rnorm(n=6, sd = sigma_det)}) |> t()

    # Make detection error covariance matrix
    Sigma_det <- diag(rep.int(sigma_det^2, times = 6))

    # Get radial error estimates for each simulated emission
    df_r <- apply(Y, 1, \(y){
      # Calculate Jacobian
      mH <- func_jacobian_x(y, receivers, c)
      # Transform detection error into positioning error
      Sigma_pos <- mH %*% Sigma_det %*% t(mH)
      # Calculate the expected value and SD of the radial error
```

```r
      lst_r <- func_radial_error(
        Sigma = Sigma_pos)# Positioning error covariance
      # Return as a data.frame
      return(as.data.frame(lst_r))}) |>
      do.call(what = 'rbind')

    # Apply TDOA positioning to each simulated emission
    x_hat <- apply(Y, 1,
      func_h,#                TDOA positioning function
      receivers = receivers,# Matrix of receiver coordinates
      c = c# m/s              Speed of signal transmission
      ) |> t()

    # Calculate the true radial error
    # (distance between the true and TDOA estimated tag location)
    r_true <- apply(x_hat, 1, \(pos) sqrt((pos - x_i) %*% (pos - x_i)))

    # Calculate the standardized residuals of the radial error
    r_stnd_resid <- (r_true - df_r$E_r) / df_r$sigma_r

    # Store results
    df_results <- rbind(
      df_results,
      data.frame(
        tag = factor(i), # Location of the tag
        sigma_det = factor(sigma_det*1000),# (ms) Detection error SD
        r_stnd_resid = r_stnd_resid))# Standardized radial error residuals
  }
}

# Plot the simulation results
lattice::bwplot(
  r_stnd_resid ~ sigma_det | tag,
  data = df_results, ylim = c(-2, 3),
  panel = \(...){
    lattice::panel.abline(h = qnorm(c(0.25,0.5,0.75)), col = 'red')
    lattice::panel.bwplot(...)
  }, pch = "|",
  ylab = "Standardized residual radial error",
  xlab = expression(paste(
    "Detection error standard deviation ",sigma["det"], " (ms)")),
  as.table = T)
```

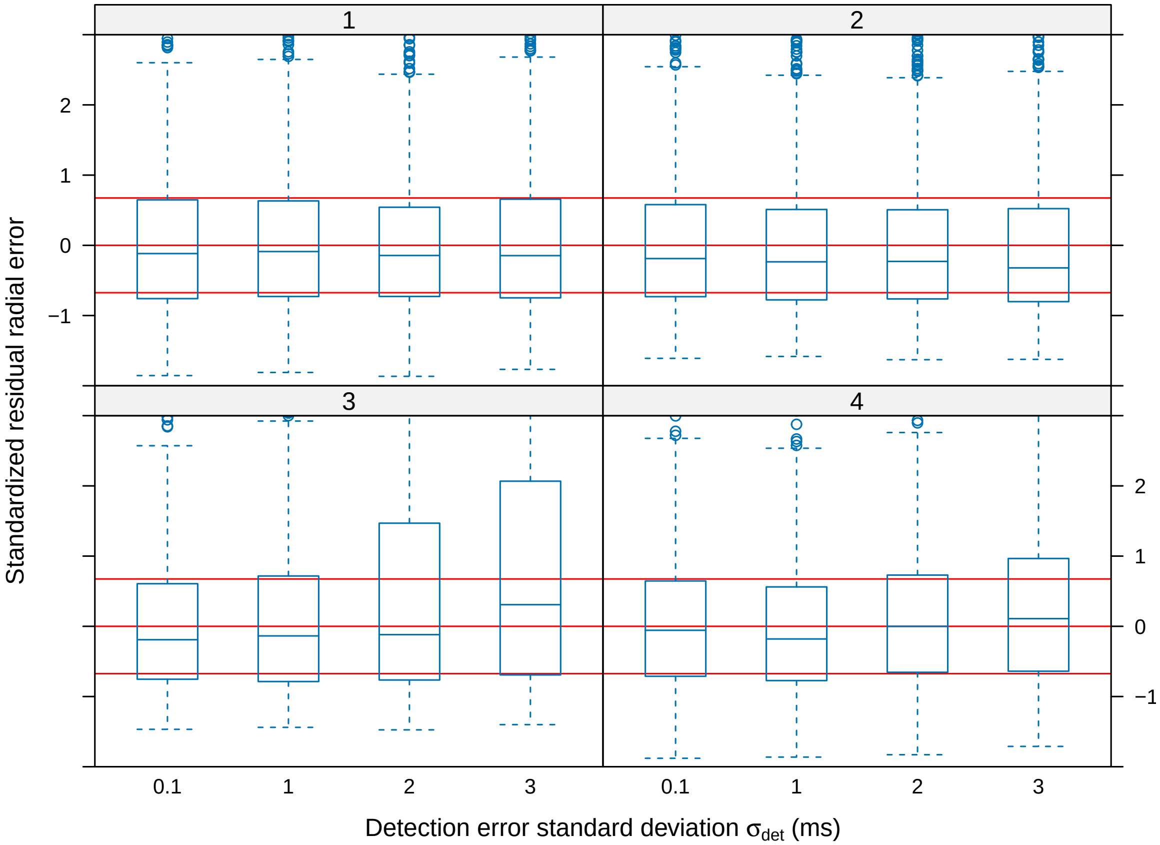


In the above plot, we can see that for the locations inside the array (1,2, and 4), the estimated residual error bears a strong resemblance to the true radial error. That is, the standardized residuals broadly follow a standard normal distribution. For location 3 which is outside the array, increasing detection error seems to result overdispersed residuals. Here, the true residual error tends to be much larger than what is reported from the estimated values, indicating that we don't place much faith in radial error estimates from this location for high detection errors ($\sigma_{\text{det}} > 1$ ms).
